# Quantifying memory CD8 T cell-mediated immune pressure on influenza A virus infection *in vivo*

**DOI:** 10.1101/2020.01.27.921031

**Authors:** Zheng-Rong Tiger Li, Veronika I. Zarnitsyna, Anice C. Lowen, Rustom Antia, Jacob E. Kohlmeier

**Affiliations:** Department of Biology, Emory University, Atlanta, Georgia, USA; Department of Microbiology and Immunology, Emory University School of Medicine, Atlanta, Georgia, USA

**Author notes:** Correspondence: Jacob Kohlmeier, 1510 Clifton Road, RRC 3133, Atlanta, GA 30322.

## Abstract

The conservation of T cell epitopes in human influenza A virus has prompted the development of T cell-inducing influenza vaccines. However, the selection pressure mediated by memory CD8 T cells upon influenza virus has not been directly measured. Using a droplet digital PCR technique to distinguish wild-type and an epitope-mutant PR8 influenza viruses *in vivo*, this study quantifies the viral replicative fitness of a CD8 T cell-escaping mutation in the immunodominant influenza NP_366-374_ epitope in C57BL/6 (B6) mice under different settings of cellular immunity. Although this mutation does not result in a viral fitness defect *in vitro* or during the early stages of *in vivo* infection in naïve B6 mice, it does confer a moderate but consistent advantage to the mutant virus following heterosubtypic challenge of HKx31-immunized mice. In addition, this advantage was maintained under increased MHC diversity but became more substantial when the breadth of epitope recognition is limited. Finally, we showed that lung-resident, but not circulating, memory CD8 T cells are the primary source of cellular immune pressure early during infection, prior to the induction of a secondary effector T cell response. Integrating the data with an established modeling framework, we show that the relatively modest immune pressure mediated by memory CD8 T cells is one of the important factors responsible for the conservation of CD8 T cell epitopes in influenza A viruses that circulate among humans. Thus, a T cell-inducing vaccine that generates lung-resident memory CD8 T cells covering a sufficient breadth of epitopes may transiently protect against severe pathology without driving the virus to rapidly evolve and escape.

**Author Summary:** Since the historic Spanish flu in 1918, influenza has caused several pandemics and become an important public health concern. The inactivated vaccines routinely used attempt to boost antibodies, which may not be as effective when antigenic mismatch happens and could drive the virus to evolve and escape due to their high immune pressure. In contrast, the ability of influenza-specific T cells to reduce pathology and the conservation of T-cell epitopes across subtypes have shed light on the development of universal vaccines. In this study, we assessed the CD8 T cell-mediated selection pressure on influenza virus in mouse using a digital PCR technique. Within mice that have influenza-specific systemic and lung-resident memory CD8 T cells established, we found the advantage conferred by an escaping mutation in one of the immunodominant epitopes is around 25%. This advantage becomes much greater when the cellular immunity focuses on the focal epitope, while it is delayed when only systemic cellular immunity is established. Combining the data with our previous modeling work, we conclude that the small selection pressure imposed by CD8 T cells can explain the overall conservation of CD8 T cell epitopes of influenza A virus in addition to functional constraint.

## Introduction

Influenza is an important public health issue, resulting in approximately 410,000 deaths worldwide annually [1]. The current inactivated influenza vaccines aim to induce antibodies that prevent viral infection; however, this strategy has two major stipulations: (i) an effective antibody response requires the match of antigenicity between circulating and vaccine strains [2, 3], and (ii) the selection pressure mediated by neutralizing antibodies may drive the virus to evolve and escape [4, 5]. Vaccines that induce broadly-neutralizing antibodies against the conserved regions of hemagglutinin (HA) and/or neuraminidase (NA), and vaccines that induce T cells targeting the conserved epitopes on the internal proteins (e.g., nucleoprotein [NP]), have been proposed to overcome these drawbacks [6–10].

Memory CD8 T cells are categorized into central (T_CM_), effector (T_EM_), and tissue-resident (T_RM_), according to their ability to circulate between secondary lymphoid organs, blood, and peripheral tissues [11–13]. Although influenza-specific memory T cells are not likely to provide sterilizing immunity, CD8 T cell responses, lung T_RM_ in particular, have been shown to prevent severe pathology in mice [14–18] and reduce symptoms in humans [19] following heterosubtypic influenza infection. Although it hasn’t been measured directly, the ability of antiviral memory CD8 T cells to limit influenza virus replication may also limit viral transmission [19, 20]. The ability of CD8 T cell to recognize and respond to diverse influenza strains and subtypes is mainly due to the conservation of CD8 T cell epitopes. Out of 64 known CD8 T cell NP epitopes across all human leukocyte antigen (HLA) alleles, only 6 epitopes have been experimentally verified to have escaping mutations [21], underpinning the potential of developing a T cell-inducing vaccine to combat influenza.

Several studies have attempted to elucidate the reason why influenza virus epitopes recognized by CD8 T cells are conserved despite cellular immune pressure. One of the mainstream hypotheses is that detrimental fitness effects would result from mutations to viral proteins that must interact with a range of host proteins to ensure efficient replication [22–24]. Although epistatic interactions with compensatory mutations may recover the viral fitness [25, 26], this process may slow the rate of generating competitive mutants. In our previous modeling work, we hypothesized that small selection pressure imposed by CD8 T cells, combined with human MHC polymorphisms, may further limit the rate of invasion of an escaping mutant [21]. Viral evolution due to CD8 T cell immunity has been observed in HIV [27, 28], HCV [29], and chronic influenza infection of immunocompromised patients [30]. However, quantitative measures of the selection pressure from memory CD8 T cells on influenza viral dynamics *in vivo* are lacking in the context of influenza infections in immunocompetent hosts.

In this study, we aim to quantify the selective advantage of a CD8 T cell-escaping influenza mutant under different contexts related to (i) distinct memory CD8 T cell subsets and (ii) the breadth of memory CD8 T cell responses. The mouse-adapted influenza strain, A/Puerto Rico/8/1934 (H1N1) (PR8), harbors three H-2*^b^*-restricted immunodominant CD8 T cell epitopes [31, 32]. We used the wild-type PR8 and PR8 with a point mutation on one of the immunodominant epitopes, NP_366-374_, such that the peptide-MHCI binding is disrupted and the mutant epitope is not presented [33]. To minimize the between-subject variation in measuring the viral kinetics, we used a droplet digital PCR system to simultaneously measure the viral loads of both wild-type and epitope-mutant viruses in a single co-infected animal at selected time points. Then, the selection coefficient of the escaping mutant was estimated by comparing the area under growth curves of the two viruses. Integrating these data with our modeling framework allowed us to quantify the factors that determine the rate of invasion of epitope-variants that escape CD8 T cell responses. Our results are consistent with the argument that broad memory CD8 T cell immunity imposes modest selection pressure on the virus, which helps explain the overall conservation of CD8 T cell epitopes of human IAV.

## Results

### *In vitro* viral growth detects no significant fitness defect of NP-N370Q mutation

Prior studies have identified a mutation in the immunodominant PR8 influenza nucleoprotein (NP)_366-374_ CD8 T cell epitope presented on H-2*^b^* that prevents loading of the peptide onto MHCI [33]. We constructed wild-type (WT) and NP-N370Q mutant (MT) PR8 influenza viruses by reverse genetics and first compared viral growth in MDCK cell culture to determine if this mutation impacted viral fitness. The viral titers at five time points after inoculation are shown in **Fig 1**, where no significant difference was detected between virus strains (three-way ANOVA, *p*-value for virus strains = 0.797). A further logistic growth model fitting found no significant difference between the estimated model parameters associated with WT and MT PR8 viruses (**Table 1**). Therefore, we conclude that the NP-N370Q mutation does not result in a fitness defect *in vitro*.

**Fig 1.**
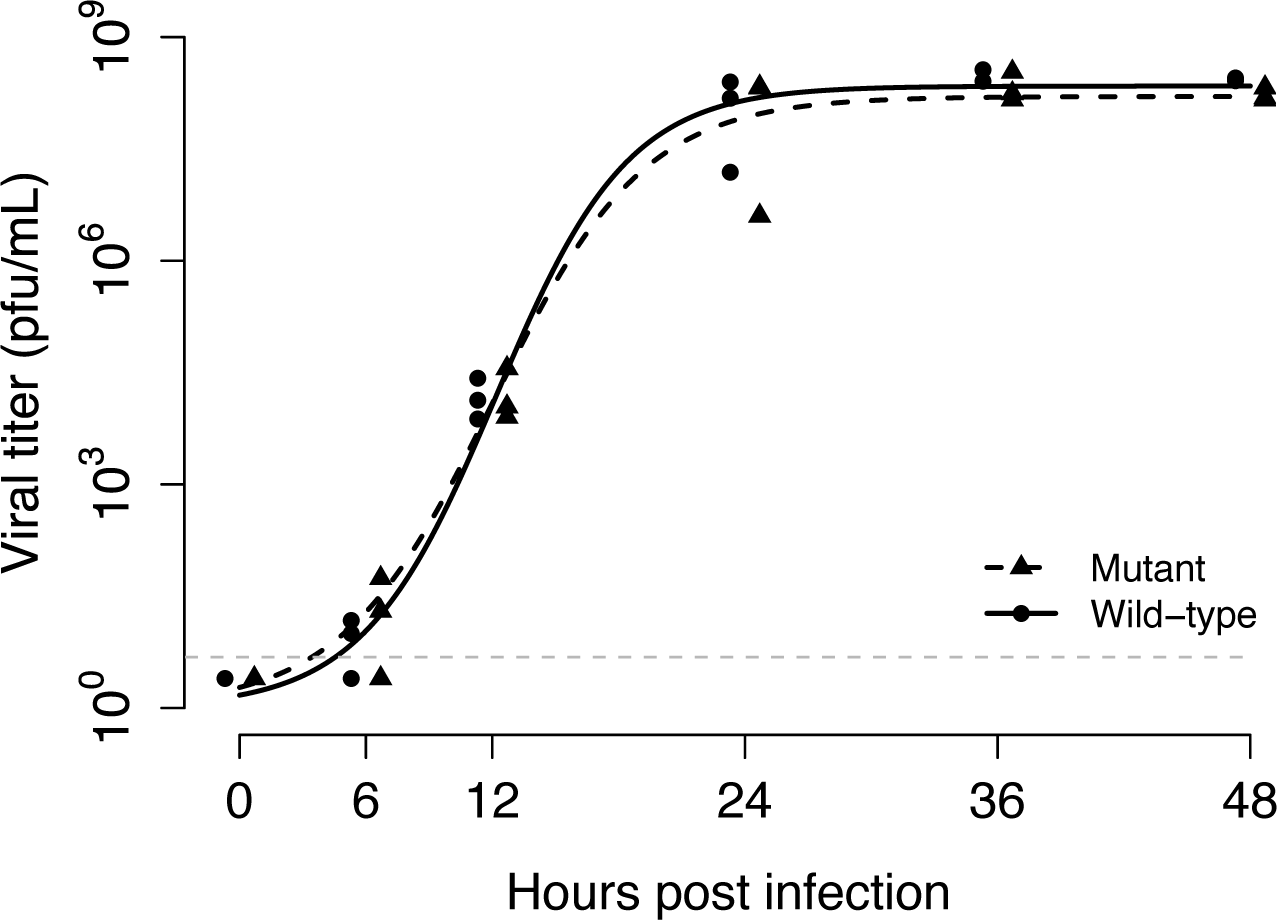
Viral growths of wild-type (WT) and NP-N370Q mutant (MT) PR8 influenza viruses in MDCK cell culture. MDCK cells were infected with WT or MT viruses at an multiplicity of infection (MOI) of 0.01. No significant difference was detected between virus strains (three-way ANOVA, *p*-value for virus strains = 0.797). The gray dotted line denotes the limit of detection (5 pfu/mL).

**Table 1.** The point estimates and 95% confidence intervals for the logistic growth model parameters

| **Table 1** | The point estimates and 95% confidence intervals for the logistic growth model parameters |  |  |
| --- | --- | --- | --- |
| Virus strain | Growth rate | Capacity | Midpoint |
| Wild-type | 0.32 [0.24, 0.43] | 8.3 [8.1, 8.6] | 12.2 [11.5, 13.0] |
| Mutant | 0.28 [0.21, 0.40] | 8.2 [7.8, 8.6] | 12.0 [11.0, 13.4] |

### Droplet digital PCR robustly differentiates between the wild type and mutant

In order to infer the impact of CD8 T cell-escaping mutations from *in vivo* experiments, we first had to design and validate a method for simultaneous measurement of WT and MT PR8 viruses within the same host. Thus, we employed a droplet digital PCR (ddPCR) system with probes specific for the WT or MT variants of the NP_366-374_ epitope (See *Supplemental Information*). The assay is highly specific for the individual strains (**Fig 2A**), as the WT probe detected positive signals only from the WT viral RNA samples, and the MT probe was similarly specific for the MT viral RNA samples. This assay also robustly reflects the change in RNA concentration, as a 10-fold dilution of input viruses resulted in a 10-fold decrease in the final readouts indicating unbiased detection of WT and MT viruses. (**Fig 2B**). Furthermore, the viral loads measured by ddPCR (in copies/mL) were consistent with viral titers measured by plaque assay (in pfu/mL), and there was no preferential detection of WT or MT viral RNA (**Table 2**). Thus, the ddPCR system allows us to accurately differentiate between WT and MT PR8 viruses in a mixed sample.

**Fig 2.**
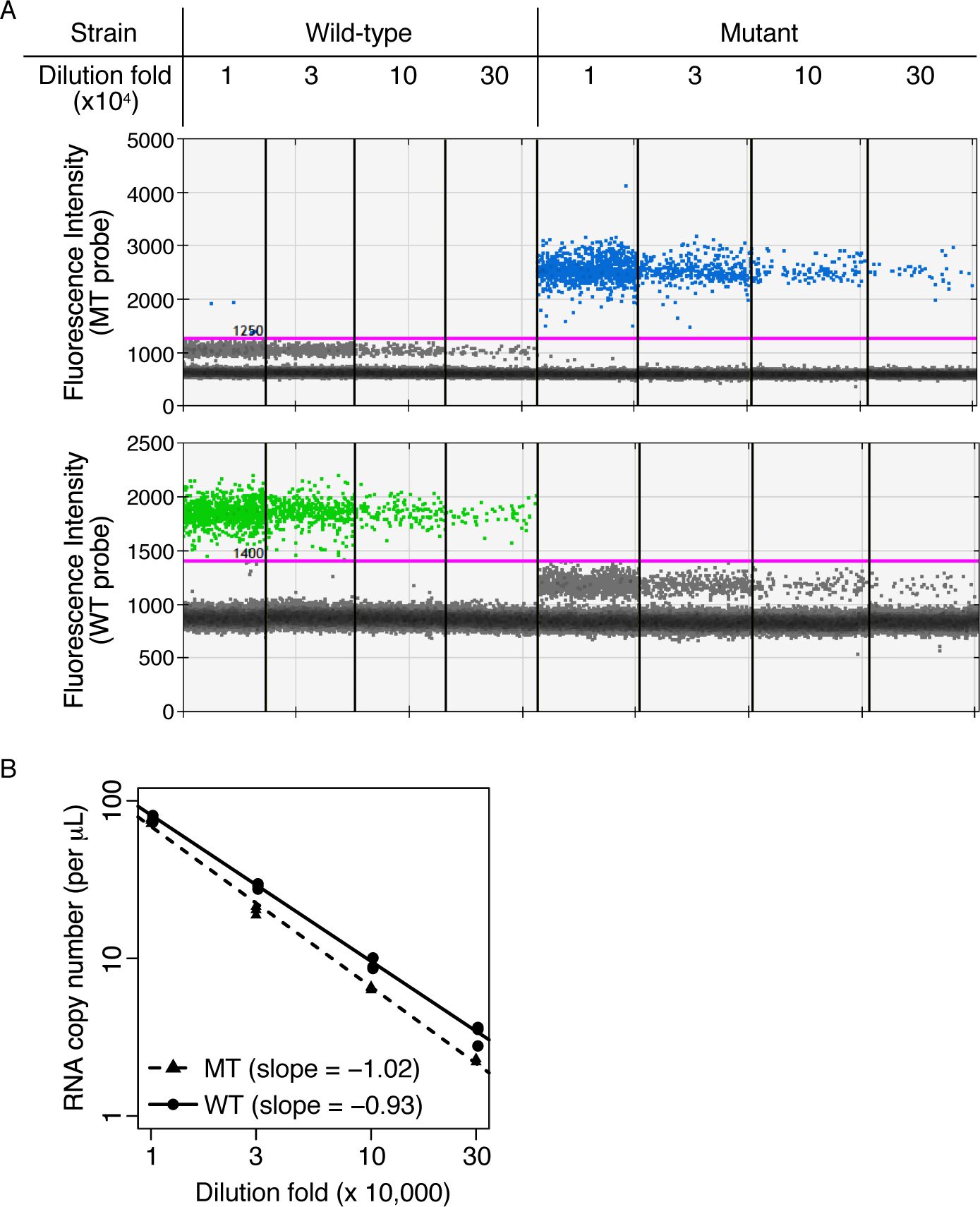
Testing the performance of droplet digital PCR (ddPCR) with serially diluted WT and MT PR8 virus stocks. (A) A representative figure of ddPCR analysis demonstrates a clear discrimination of positive signals detected by the MT probe (blue dots in the top panel) or the WT probe (green dots in the bottom panel) from negative signals (gray dots in both panels). (B) Regression lines quantified from the data in (A) over a range of virus dilutions.

**Table 2.** Comparison of viral load measured by droplet digital PCR (copy number/mL) and viral titer measured by plaque assay (pfu/mL).

|  |  |  |  |
| --- | --- | --- | --- |
| **Table 2** | Comparison of viral load measured by droplet digital PCR (copy number/mL) and viral titer measured by plaque assay (pfu/mL). |  |  |
| Virus strain | RNA copy number<br>(10<sup>8</sup> per mL) | Plaque forming unit<br>(10<sup>8</sup> per mL) | Ratio<br>(RNA copy / pfu) |
| Wild-type | 85 (± 6.0) | 10.5 (± 1.5) | 8.1 |
| Mutant | 67 (± 8.3) | 9.8 (± 1.5) | 6.9 |

### Wild type and mutant viruses have similar fitness during the early stage of primary infection

To determine if the MT PR8 virus has any fitness defect *in vivo* in the absence of influenza-specific memory CD8 T cell immunity, we infected naïve C57BL/6 (B6) mice with an equal mixture of WT and MT PR8 viruses (**Fig 3A**). In naïve B6 mice, the viral loads of WT and MT viruses in the lungs recapitulated the kinetics reported previously [34], where they (i) increased exponentially and peaked around 3 days post-infection (dpi), (ii) slowly decayed from 3 to 5 dpi, and (iii) quickly decayed and were cleared after 5 dpi (**Fig 3B**). Across the 5 time points being sampled, only on day 5 was the average log-transformed viral load of MT significantly higher than the WT (*p* = 0.0007).

**Fig 3.**
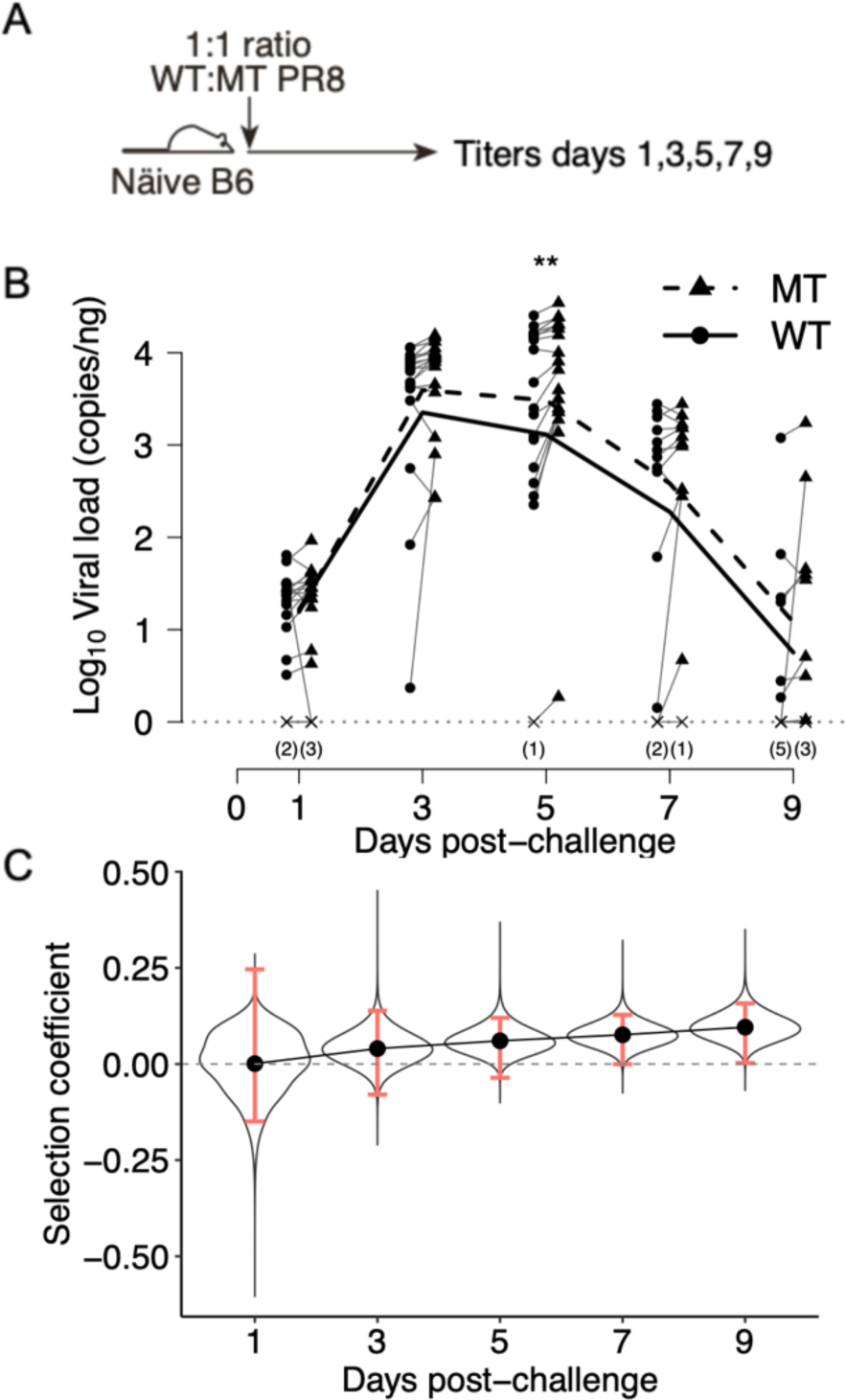
Comparison of the wild-type (WT) and mutant (MT) kinetics in primary infection. (A) Naïve C57BL/6 (B6) mice were inoculated with an equal mixture of WT and MT PR8 viruses, and the viral loads were measured on selected time points. (B) *In vivo* viral kinetics on days 1, 3, 5, 7, and 9 post-challenge as measured by ddPCR. Detectable WT, detectable MT, and undetectable viral loads are marked by circles, triangles, and crosses, respectively. Paired data points are connected with a line. Numbers of undetectable data points are indicated in the parentheses. (C) The selection coefficient of MT and sampling distributions measured by AUC and bootstrapping. The point estimates and 99% CIs for the selection coefficient of MT are indicated by the dots and the red bars, respectively.

We then defined the *selection coefficient of MT* as

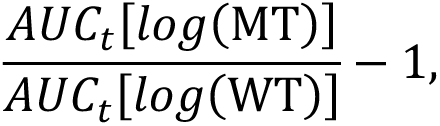

where AUC*_t_* stands for the area under viral growth curve from days 0 to *t*. This statistic quantifies the fitness change associated with MT as a fraction of the fitness of WT. For instance, 0.1 indicates the MT has 10% higher fitness compared to WT, while −0.15 indicates the MT has 15% lower fitness compared to the WT. The sampling distribution of this statistic was assessed by the bootstrapping method (See *Supplemental Information*).

The selection coefficient of MT gradually increased from 0.001 on day 1 to 0.1 on day 9, while the 99% empirical bootstrap confidence interval (99% CI hereafter) did not contain zero only on day 9 (**Fig 3C**). These findings showed that the MT virus does not have a detectable fitness defect during the early stages of infection *in vivo*, but gains an advantage as the infection progresses, likely due to its ability to escape from the immunodominant NP_366-374_-specific effector CD8 T cells, similar to previous reports [34].

### The mutant virus acquires selective advantage early during secondary heterosubtypic infection in the presence of influenza-specific memory CD8 T cell immunity

To estimate the selective advantage of the MT virus under different settings of cellular immunity, we intranasally (i.n.) primed B6 mice with HKx31 (H3N2), challenged the mice 30 days later with an equal mixture of WT and MT PR8 (H1N1) viruses, and measured the viral kinetics (**Fig 4A**). Viral loads peaked on day 2, slowly decayed from days 2 to 4, and rapidly decayed and cleared from days 4 to 8. Despite a similar overall pattern to naïve mice, the peak viral loads were around 10-fold lower, and clearance was faster. The means of log-transformed MT viral loads were significantly higher than those of WT on days 1 and 4 (*p* = 0.034 and 0.0018, respectively) (**Fig 4B**). Likewise, the selection coefficient of MT continuously increased from 0.15 on day 1 to 0.27 on day 9, while the 99% CIs contained zero only on day 1 (**Fig 4C**). Thus, the MT acquired an advantage in the early stage of infection, and the advantage persisted and increased through the infection course.

**Fig 4.**
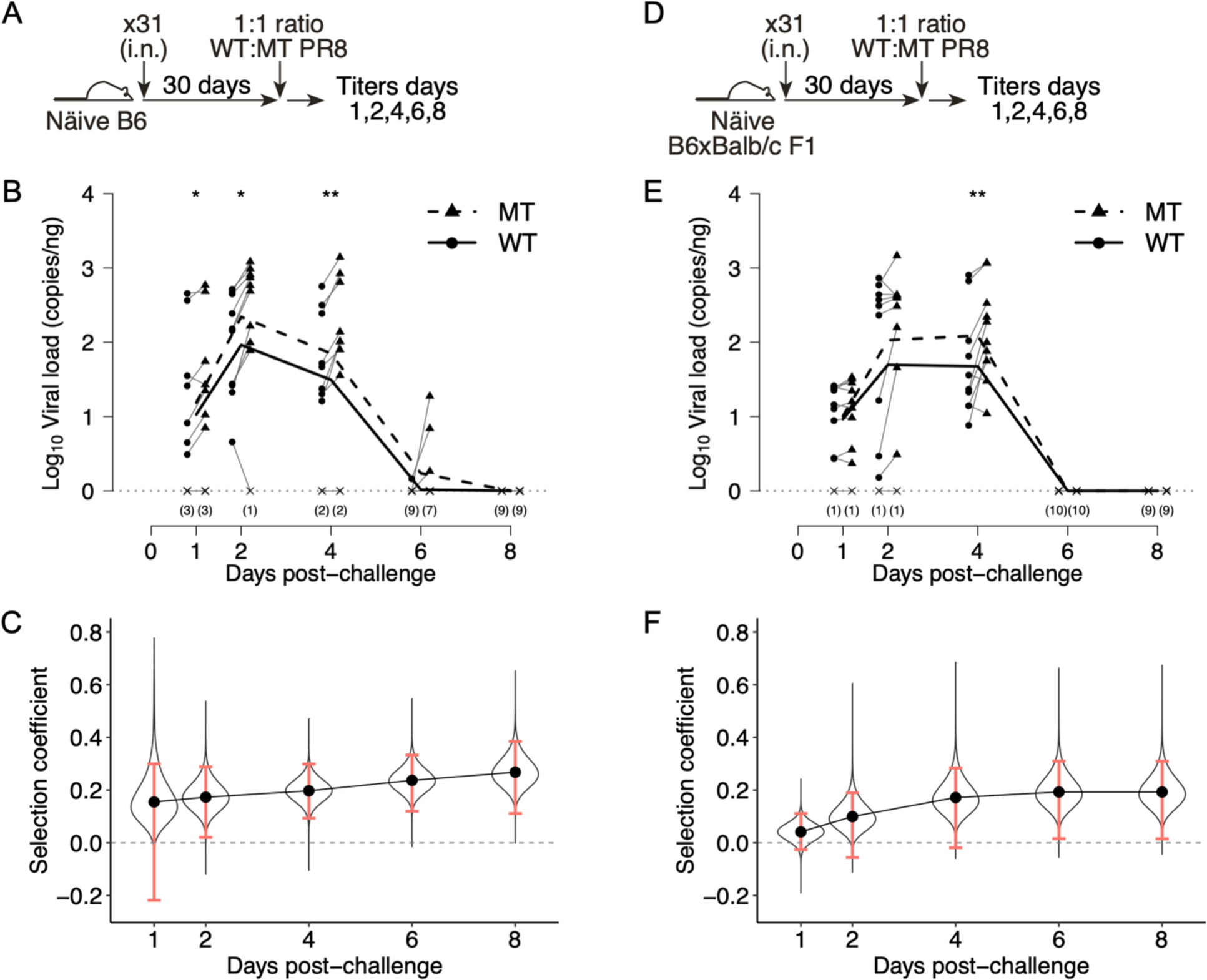
Viral kinetics in intranasally (i.n.) HKx31 (x31)-primed B6 (H-2*^b/b^*) and CB6F1 (H-2*^d/b^*) mice. (A) B6 mice were immunized i.n. with x31 and challenged with an equal mixture of WT and MT PR8 viruses 30 days later. (B) *In vivo* viral kinetics of i.n. x31-primed B6 mice on days 1, 2, 4, 6, and 8 post-challenge as measured by ddPCR. (C) The selection coefficient of MT in i.n. x31-primed B6 mice as measured AUC and bootstrapping. The 99% CIs did not contain zero except on day 1. (D) CB6F1 (F1) mice, which are the heterozygotes of H-2*^b^* and H-2*^d^* haplotypes, were immunized i.n. with x31 and challenged with an equal mixture of WT and MT PR8 viruses 30 days later. (E) *In vivo* viral kinetics of i.n. x31-primed F1 mice on days 1, 2, 4, 6, and 8 post-challenge as measured by ddPCR. (F) The selection coefficient of MT and sampling distribution in i.n. x31-primed F1 mice as measured by AUC and bootstrapping.

To investigate whether an increased MHC diversity impacts the selective advantage of the MT virus, we applied the same prime-challenge procedure to CB6F1 (F1) mice, which are the offspring of C57BL/6 and BALB/c mouse strains, and harbor both H-2*^d^* and H-2*^b^* haplotypes (**Fig 4D**). Thus, F1 mice develop a broader influenza-specific CD8 T cell response that encompasses epitopes presented by both MHC alleles. We observed similar viral kinetics in F1 mice compared to B6 mice; however, the viruses were cleared even faster than B6 mice (no virus was detected on days 6 and 8), and the MT significantly outgrew WT only on day 4 (*p* = 0.0054) (**Fig 4E**).

When looking at the selection coefficient of the MT virus, we noticed two interesting differences between i.n.-primed B6 and F1 mice (**Fig 4F**). First, the point estimates were much lower on the first two days in F1 than B6 mice (day 1: 0.04 in F1 vs. 0.15 in B6; day 3: 0.1 in F1 vs. 0.17 in B6), and the 99% CIs contained zero on both days; however, this deviation became negligible on day 4. Second, after day 4, the selection coefficient of MT showed a pronounced plateau in F1 mice, corresponding to the fact that both WT and MT viruses were cleared on days 6 and 8. Together, these data suggest that escape from cellular immunity specific for the NP_366-374_ epitope confers a fitness advantage to the MT virus early during heterosubtypic influenza infection. The impact from increased breadth of CD8 T cell response was not constant across the infection course; instead, it depends on the time of observation.

### CD8 T cell-escaping mutations confer a much greater advantage to the MT virus when the breadth of CD8 T cell response is limited

Intranasal priming of B6 mice with x31 generates memory CD8 T cells specific for multiple epitopes in addition to the immunodominant NP_366-374_ epitope, including two additional immunodominant and at least eight subdominant epitopes [31, 32]. Therefore, even if the MT PR8 virus escapes detection from NP_366-374_-specific CD8 T cells, it remains subject to recognition by CD8 T cells targeting other epitopes. However, some vaccine approaches are designed to focus the CD8 T cell response to a few immunodominant epitopes, raising the question of the selective advantage that can be gained by a mutant influenza virus that can escape a focused influenza-specific memory CD8 T cell repertoire. To address this question, we immunized B6 mice with a recombinant, replication-deficient adenovirus 5 expressing the PR8 influenza nucleoprotein (AdNP), which generates memory CD8 T cells specific for only NP-derived epitopes, mainly the immunodominant NP_366-374_ epitope [35], and challenged them 30 days later with a mixture of WT and MT PR8 viruses (**Fig 5A**). Vastly different growth kinetics of the MT virus were observed compared to the WT virus (**Fig 5B**). The MT grew continuously and peaked around day 4, at a viral load 35-fold higher than WT. After day 4, both viruses decayed at the same rate exponentially, but neither were completely cleared on day 8. At each time point selected, the means of log-transformed MT viral loads were significantly higher than those of WT (*p* = 0.031, 0.0004, 0.0002, 0.0016, and 0.023 for days 1, 2, 4, 6, and 8, respectively). Consistent with this, the selection coefficient of MT continuously climbed throughout the infectious course from 0.24 on day 1 to 0.8 on day 8, and all the 99% CIs did not contain zero (**Fig 5C**). These data show that when the antiviral memory CD8 T cell repertoire is comprised of only a single epitope, the selective advantage conferred by the corresponding escaping mutation is substantially high, much greater than in the context where memory CD8 T cells recognize multiple epitopes.

**Fig 5.**
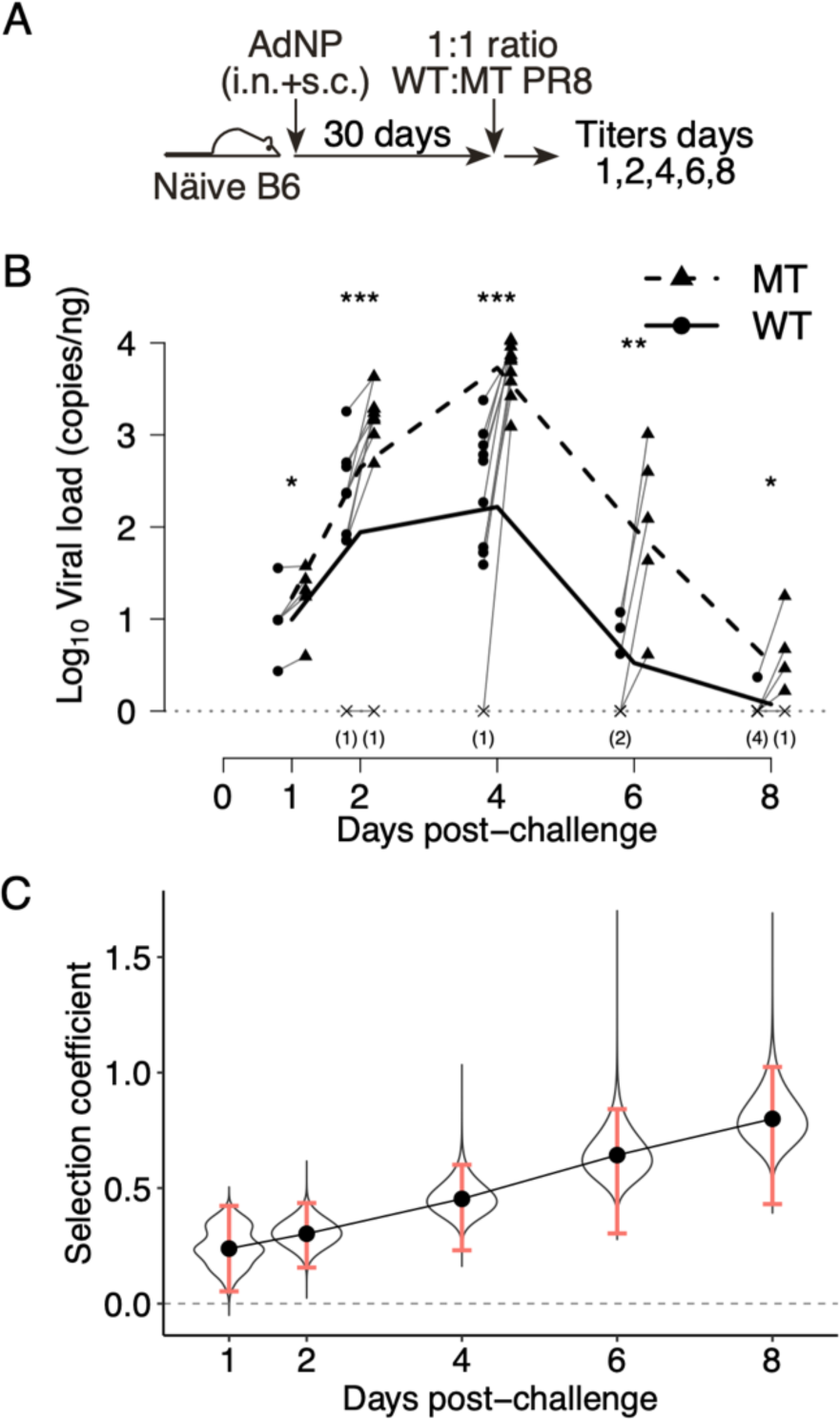
Viral kinetics in the AdNP-primed B6 mice, where only the NP_366-374_-specific memory CD8 T cells were present. (A) B6 mice were i.n. and subcutaneously (s.c.) immunized with AdNP and challenged with an equal mixture of WT and MT PR8 viruses 30 days later. (B) *In vivo* viral kinetics of AdNP-primed B6 mice on days 1, 2, 4, 6, and 8 post-challenge as measured by ddPCR. (C) The selection coefficient of MT and sampling distributions in AdNP-primed B6 mice as measured by AUC and bootstrapping. All the 99% CIs did not contain zero.

### Lung-resident memory CD8 T cells are the primary source of selection pressure during the early stage of secondary infection

Lung-resident memory CD8 T cells (lung T_RM_) are important for mediating heterosubtypic immunity and controlling influenza virus replication due to their localization in the respiratory tract enabling rapid detection of infected cells. Thus, we hypothesize lung T_RM_ impose much of the early cellular immune pressure on the virus. We tested this hypothesis by comparing the viral kinetics in 30-day i.n. x31-primed B6 mice with 30-day intramuscularly (i.m.) x31-primed B6 mice (**Fig 6A**), which we previously showed to have approximately the same number of influenza-specific systemic memory CD8 T cells as i.n.-primed mice but lack lung T_RM_ [17]. We observed that, in i.m. x31-primed mice, (i) the viral loads peaked at a later time point (day 4) and were about 6- and 12-fold higher than the peaks seen in i.n. x31-primed mice, (ii) the MT and WT viruses grew at the same rate between days 1-4 and peaked at similar levels, and (iii) after day 4, the WT virus was cleared faster than the MT virus (*p* = 0.021 on day 6, *p* = 0.0095 on day 8) (**Fig 6B**). Interestingly, the selection coefficient of MT showed a U-shape trend; it decreased from 0.12 on day 1 to 0.057 on day 4, and then increased to 0.22 on day 8 (**Fig 6C**). Nevertheless, these data showed the MT virus does not acquire the same increased advantage during the early stage of infection when the lung T_RM_ is absent. Thus, the selection pressure mediated by lung T_RM_ is the main driver of the outgrowth of MT virus observed in i.n. x31-primed B6 mice.

**Fig 6.**
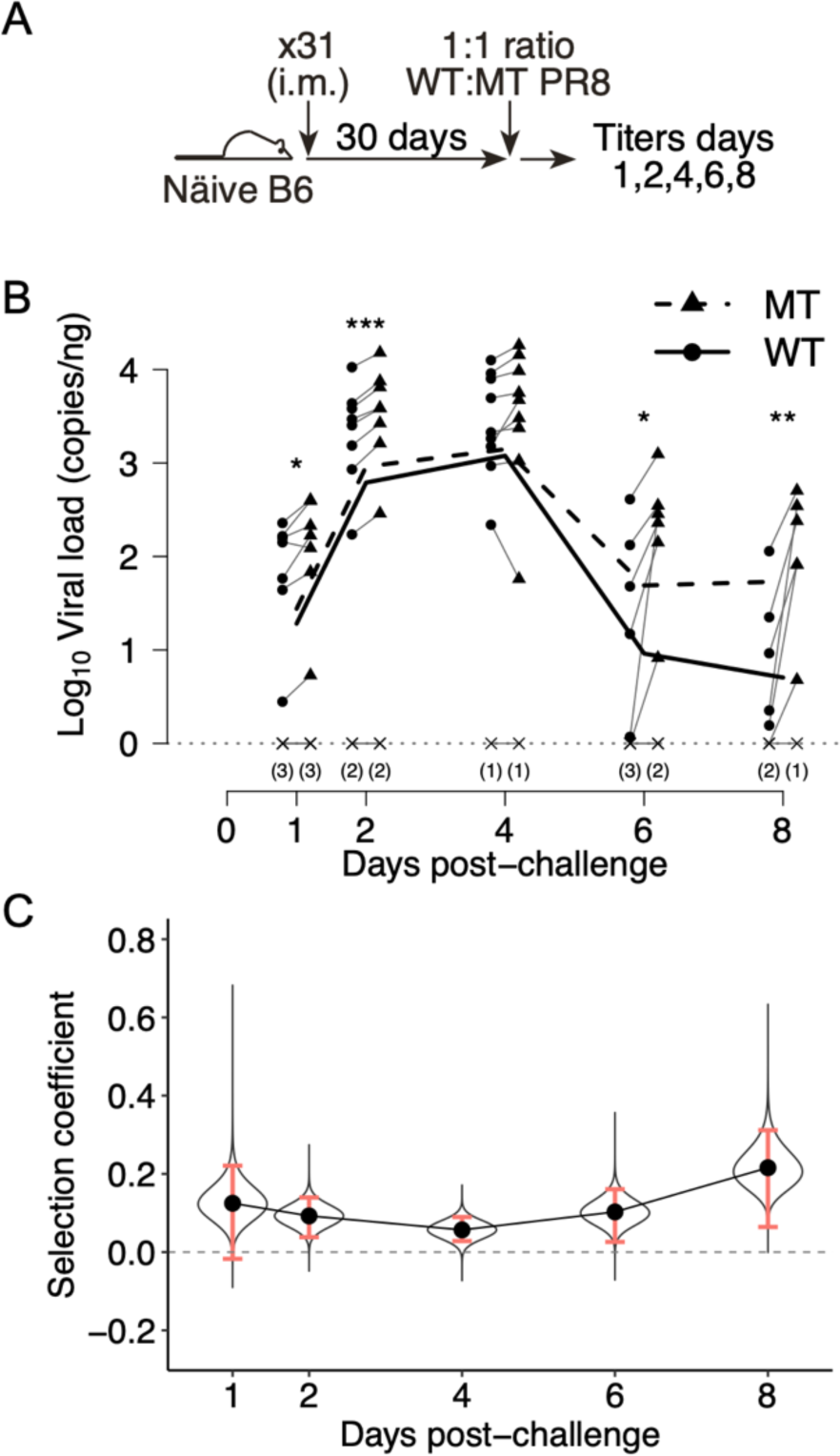
Viral kinetics in intramuscularly (i.m.) x31-primed B6 mice that lack lung T_RM_. (A) B6 mice were i.m. immunized with x31 and challenged with an equal mixture of WT and MT PR8 viruses 30 days later. (B) *In vivo* viral kinetics of i.m. x31-primed B6 mice on days 1, 2, 4, 6, and 8 post-challenge as measured by ddPCR. (C) The selection coefficient of MT and sampling distributions in i.m. x31-primed B6 mice as measured by AUC and bootstrapping. The 99% CIs did not contain zero except day 1.

### Comparison of the selection coefficients of MT virus revealed how cellular immunity impacts the selective advantage acquired by escaping mutations

We summarized the selection coefficients of MT virus estimated from log-transformed viral kinetics in **Table 3**. Overall, the MT had higher fitness than the WT through the infection course, regardless of the context of pre-existing cellular immunity. The fitness gain ranged from 10% to 74%, depending on the immune settings. However, a stratifying analysis revealed that the fitness gain does not evenly distribute across the infection course. During the first 4 or 5 days, the MT had little advantage in naïve and i.m. x31-primed mice. This implied most of the advantage was acquired later during infection under these scenarios. In contrast, the MT acquired around 15% increase in fitness in both i.n. x31-primed B6 and F1 mice through days 0-4, but over the whole infection course this advantage increased to 24% in B6 mice while it was maintained in F1, corresponding to faster viral clearance in F1 mice. Lastly, the MT acquired a large and stable fitness gain in AdNP-primed B6 mice, at an average rate 8% per day. In summary, the NP-N370Q mutation confers no more than 25% increase in fitness when CD8 T cell immunity targets multiple epitopes, while it can confer up to 74% increase in fitness when only the NP_366-374_ epitope is targeted.

**Table 3.** Point estimates and 99% empirical bootstrap confidence intervals for the selection coefficient of MT based on log-transformed viral kinetics.

| **Table 3** | Point estimates and 99% empirical bootstrap confidence intervals for the selection coefficient of MT based on log-transformed viral kinetics. |  |  |  |
| --- | --- | --- | --- | --- |
|  | Days 0-4 |  | Days 0-8 |  |
| Naïve B6<sup>†</sup> | 0.06 | [-0.036, 0.12 ] | 0.096 | [ 0.003, 0.16 ] |
| i.n. x31 B6 | 0.20 | [ 0.093, 0.30 ] | 0.27 | [ 0.11 , 0.38 ] |
| i.n. x31 F1 | 0.17 | [-0.019, 0.28 ] | 0.19 | [ 0.015, 0.31 ] |
| AdNP B6 | 0.45 | [ 0.23 , 0.60 ] | 0.80 | [ 0.43 , 1.02 ] |
| i.m. x31 B6 | 0.057 | [ 0.028, 0.09 ] | 0.22 | [ 0.064, 0.31 ] |
| <sup>†</sup> For naïve mice, point estimates and bootstrap were conducted over days 0-5 and days 0-9. |  |  |  |  |

We explored the reason why escape from NP_366-374_-specific CD8 T cells only confers modest selective advantage, focusing on CD8 T cell kinetics during secondary infection. We challenged i.n. x31-primed B6 mice with either WT or MT viruses alone and tracked the numbers and percentages of CD44^+^Tetramer^+^ CD8 T cells in the lung interstitium on days 0, 2, 4, and 7 (**Fig S1A**). The percentage of D*^b^*NP_366_-specific CD8 T cells increased from 5% to around 50% in the WT-challenged mice, but shrank to 1.5% in the MT-challenged mice. In contrast, the D*^b^*PA_224_- and K*^b^*PB1_703_-specific CD8 T cells increased from 14% to 40% in the MT-challenged mice, distinct from the modest increase of 10% to 13% observed in the WT-challenged mice. The kinetics in the airways had greater variation but followed the same pattern (**Fig S1B**). These data suggest the loss of D*^b^*NP_366_-specific CD8 T cell response might be compensated by the CD8 T cells against other epitopes.

Finally, we attempted to infer the transmission fitness of the viruses based on their replicative fitness, which is estimated by the AUC of viral kinetics. Since the relationship between transmissibility and viral loads in the respiratory tract hasn’t been well established, we conducted the AUC and bootstrap analysis assuming the relationship is linear or sigmoidal (**Table S1**. Also see **Fig S2** for the sensitivity analysis on the choice of parameters of Hill function). Although different point estimates and 99% CIs were obtained, the choice of link function did not qualitatively change the inference about CD8 T cell-mediated selection pressure.

## Discussion

T cell-inducing vaccines have been proposed as a potential means to lessen the burden of influenza disease due to their ability to limit viral replication and immunopathology and the high conservation of immunodominant CD8 T cell across different influenza strains [6–10]. Whether these vaccines may drive the evolution of cellular immune escape influenza variants has not been investigated comprehensively, partly due to the lack of empirical measures for selection pressure mediated by memory CD8 T cells. In this study, we measured the replicative fitness of wild-type PR8 and an immunodominant CD8 T cell-escaping mutant, PR8 NP-N370Q, in mice under different settings of cellular immunity, and we infer the epidemiological fitness of the CD8 T cell-escaping mutant by integrating the data with established modeling frameworks.

The selection coefficient of the mutant virus compared to the wild-type virus measures fitness change associated with the NP-N370Q mutation. Consistent with *in vitro* viral growth characteristics, the mutant virus does not show a significant fitness defect in the absence of pre-existing cellular immunity during the early stage of infection, as demonstrated by *in vivo* fitness measures in naïve B6 mice. During the late stage of infection, the MT virus acquired a slight advantage (around 10%), likely due to the increasing selection pressure of newly-induced effector NP_366-374_-specific CD8 T cells that recognize only the WT virus [34]. The selective advantage became significantly larger within intranasally x31-primed mice (16-24% in B6 and 15-17% in F1), where the selection pressure of NP_366-374_-specific memory CD8 T cells is present from the onset of infection. Summarizing the data from these scenarios showed the MT virus has a 15-25% advantage in replication by escaping the immunodominant NP_366-374_-specific memory CD8 T cell response, much smaller than the advantage it would acquire by escaping the neutralizing antibodies, which block viral entry and thus prevent replication all together.

Why is the selective advantage conferred by escaping the NP_366-374_-specific memory CD8 T cells relatively small? A likely mechanism is that the ability to evade the NP_366-374_-specific response is compensated by memory CD8 T cells specific for other immunodominant epitopes, namely, the PA_224-233_ and PB1_703-711_ epitopes [36, 37]. This compensation was evident in B6 mice primed with x31 and challenged with either WT or MT viruses alone, where expansion CD8 T cells specific for the additional immunodominant FluPA and FluPB1 epitopes was evident only in mice challenged with MT virus (**Fig S1**). Overall, these data suggest that the breadth of epitope recognition by CD8 T cells enables the cellular immune response to at least partially compensate for the loss of a single immunodominant epitope. However, previous studies have shown the presentation of epitopes vary across different cell types [38], and that may also impact the advantage a virus may gain through an escaping mutation.

Interestingly, in the MHC heterozygote (H-2*^b/d^*) F1 mice, the advantage acquired by the MT virus started with a lower value, reached the same level as B6 mice on day 4, and leveled off thereafter. We conjectured that this difference stems from two mechanisms. First, NP_366-374_-specific CD8 T cells contribute to a majority of the cellular immune response to H-2*^b^*-restricted epitopes in the lungs during a secondary influenza infection [36, 37], and its loss may not be fully compensated by the additional H-2*^d^*-restricted CD8 T cells. This results in the similarity between B6 and F1 on day 4 after PR8 challenge. Second, the cellular immune response can be affected by the differential presentation of epitopes by distinct cell types. For instance, the PA_224-233_ is highly expressed by dendritic cells but weakly expressed by epithelial cells, resulting in delayed viral clearance after influenza infection [38]. Potentially, the H-2*^d^*-restricted CD8 T cells may have different dynamics compared to the NP_366-374_-specific CD8 T cells. This results in the difference before day 2 and after day 6. Studies elucidating the dynamics of H-2*^d^*-restricted CD8 T cells in the BALB/c and CB6F1 mice would be needed to test this argument. There is also a possibility that heterozygosity of immune-related genes in F1 mice has a broader impact on antiviral immunity beyond MHC restriction, although this would be anticipated to affect both the WT and MT viruses equally.

Given the strong interest in designing vaccines that can promote effective cellular immunity against influenza, it is important to understand how different scenarios of T cell memory can impact the selective advantage of CD8 T cell-escaping mutations. Thus, we directly assessed the impact of two practical scenarios on viral fitness: a lack of lung-resident CD8 T cell memory, and a memory CD8 T cell response focused on a single epitope. Intramuscular (i.m.) injection is the conventional influenza vaccination route in humans; however, previous studies have shown the i.m. immunization route does not induce lung T_RM_, which are essential for optimal cellular immunity against heterologous influenza challenge [14–18]. Our data reveal that lung CD8 T_RM_ are the primary source of selection pressure during the early stage of influenza virus infection, as the MT virus outgrew the WT virus in the i.n. x31-primed mice at all times over the course of infection, but failed to do so until day 6 post-challenge in the i.m. x31-primed mice. In contrast, when the influenza-specific memory CD8 T cell pool is directed against a single epitope as seen in AdNP-immunized mice, the mutant acquired a substantially large advantage during both the early stage (40%) or across the whole infection course (74%). These two cases demonstrate that, while inducing a population of lung CD8 T_RM_ specific for one single epitope may be effective in controlling early viral replication [35], it would also impose a strong selection pressure on the virus.

This study sheds light on the design of T cell-inducing vaccines and the consequences of vaccination. Given the assumption that transmission fitness is proportional to log-transformed replicative fitness, the advantage of one CD8 T cell-escaping mutation is limited to under 25% when the infections occur through the natural route (i.e., airborne transmission), even if the mutant escapes from immunodominant CD8 T cells in MHC-homozygous individuals. This indicates that a vaccine inducing lung CD8 T_RM_ of sufficient breadth to recognize multiple epitopes could provide effective protection by controlling early viral replication and lessening immunopathology with limited selection pressure against a single epitope. Additionally, to our best knowledge, the protection mediated by lung CD8 T_RM_ is transient. In mice, the lung CD8 T_RM_ immunity wanes within 6 to 7 months after initial infection [16, 39, 40]. Although the waning time in human remains unclear, as people getting influenza infection every 5 to 7 years [41], the selection pressure from lung CD8 T_RM_ at individual level would be much smaller than what we observed in this study. Combining the data with our prior modeling work [21], we suggested the conservation of CD8 T cell epitopes in influenza A viruses can be explained by the small and transient selection pressure from cellular immunity as well as the potential fitness defect due to escaping mutations.

It stands to reason that viral replication and viral transmission are linked, but exact function linking these two quantities has not been well established. As different link functions may end up with different estimates of transmission fitness, this limits the quantitative inference from integrating experimental data with the model. Although there are robust animal transmission models of influenza available, such as ferrets and guinea pigs, their influenza-specific CD8 T cell profiles are not well characterized; in contrast, mice are an excellent model to investigate T cell immunity but they do not transmit influenza virus. By running a sensitivity analysis based on Hill function, we show that the choice of parameters does change the estimates of selection coefficient and, thus, the quantitative inference of the time required for a CD8 T cell-escaping mutant virus to invade the circulating viral population. These findings highlight the need for development of an influenza transmission animal model with well-characterized influenza-specific T cell profiles for linking the measurements of replicative, transmission, and epidemiologic viral fitness.

## Materials and Methods

### Viruses

The HKx31 (H3N2) virus was amplified in 10- to 11-day-old embryonated chicken eggs as previously described [17]. The wild-type (WT) PR8 (H1N1) virus and the NP-N370Q mutant (MT) virus were generated using the reverse genetics system [42].

### Mice and infections

C57BL/6 (H-2*^b/b^*) and CB6F1 (H-2*^b/d^*) mice were purchased from Jackson Laboratory and inoculated at 8 weeks of age. For primary infection, mice were inoculated with 50 uL of 1:1 mixture of WT and MT PR8, 125 PFU for each (total dose is 250 PFU, corresponding to 1 LD50 of PR8). For secondary infection, mice were primed with (i) 3×10^4^ EID50 of HKx31 in 30 uL HBSS intranasally (i.n.), (ii) 10^6^ EID50 of HKx31 in 50 uL HBSS intramuscularly (i.m.), or (iii) AdNP virus in 30 uL of PBS both i.n. and subcutaneously (s.c.) [35]. Thirty days later, the immunized mice received a 50 uL of 1:1 mixture that consists of 1,250 PFU of each PR8 virus (total dose is 2,500 PFU, corresponding to 10 LD50). Challenged mice were euthanized with Avertin on the indicated days post-infection. All mouse studies were approved by the Institutional Animal Care and Use Committee of Emory University.

### Measuring viral load using droplet digital PCR

RNA of virus stocks was isolated using QIAamp Viral RNA Mini Kit and stored at −80C until use. For RNA isolation from infected mice, lungs were minced and preserved in RNALater at −80C until all the samples of the same batch were collected. Total RNA was isolated from lung homogenates using Ambion RNA Isolation Kit and the concentration was measured by Nanodrop. For reverse transcription and ddPCR, cDNA from the total RNA of lungs or viral RNA was made using Maxima RT and universal influenza primers. The cDNA samples then underwent 10-fold serial dilution and were quantified by ddPCR with probes designed to discriminate wild-type and NP_366-374_ mutant epitopes. The RNA copy number data were back-calculated to the RNA concentration (copy number per ng of total RNA, see *Supplemental Information*).

### Quantitative analysis

#### Statistical tests

The *in vitro* viral titer data were tested by a three-way ANOVA to account for (i) the virus strain effect, (ii) temporal effect, and (iii) batch effect. The data were then used to estimate the parameters and the 95% confidence intervals of the logistic growth model. The log-transformed *in vivo* viral load data were tested by Student’s *t*-test for paired data. All the analyses were done in R 3.6.1.

#### Area under the curve and bootstrapping

The area under the *in vivo* viral growth curve (AUC) represents the total amount of virus produced over the course of infection and serves as a marker for the replicative fitness. Based on our assumption that the transmission fitness is related to replicative fitness, we log-transformed the viral load data and calculated the AUCs of the mutant and the wild type viruses. Bootstrapping was performed one million times to assess the uncertainty of estimation (see *Supplemental Information*). The 99% empirical bootstrap confidence intervals were calculated following basic bootstrap method [43].

## Acknowledgements

This project was supported by NIH grants R01HL122559 (J.E.K.), U01HL139483 (J.E.K. and V.I.Z.), and U19AI117891 (R.A.).

## Supplemental Information

### Maxima RT

Mix 4 uL of 5x RT buffer, 1 uL of universal F(A) primer (6 uM), 1 uL of dNTPs (10 mM), 1 uL of ribolock RI, 1 uL of Maxima RT with either 12 uL of viral RNA isolated from virus stock or 2 to 4 ug of total RNA isolated from the lungs on ice. Run for 30 minutes at 55C, 10 minutes at 85C, and hold at 10C.

### Droplet Digital PCR

A mixture containing 11 uL of 2x ddPCR supermix for probes, 1.1 uL of mixed forward and reverse primers (36 uM), 1.1 uL of wild-type probe, 1.1 uL of mutant probe, 3.3 uL of ultrapure water, and 4.4 uL of cDNA sample was used to amplify a fragment containing the WT or MT NP_366-374_ epitope. 20 uL of this mixture was added to 70 uL of droplet generation oil, and after the droplet generation step, 40 uL of the suspension was used to perform ddPCR in a 96-well PCR plate. The PCR steps are listed below:

| Temperature (C) | Time |  |
| --- | --- | --- |
| 95 | 10 mins |  |
| 94 | 30 secs | 35-40 cycles |
| 60 | 1 min |  |
| 98 | 10 mins |  |
| 4 | (Hold) |  |

The fluorescent signal was read by a QX200 Droplet Reader (BioRad) and analyzed with QuantaSoft software. The gating for positive droplets was set according to the positive and negative controls on each plate. The viral loads (*V*, copy number/ng of RNA) were calculated by the formula

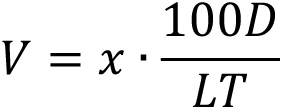

where x is the readout of ddPCR (copy number/uL), *D* is the dilution factor, *L* is the volume of RNA used for RT (uL), *T* is the RNA concentration measured by Nanodrop (ng/uL), and 100 is a coefficient with unit uL.

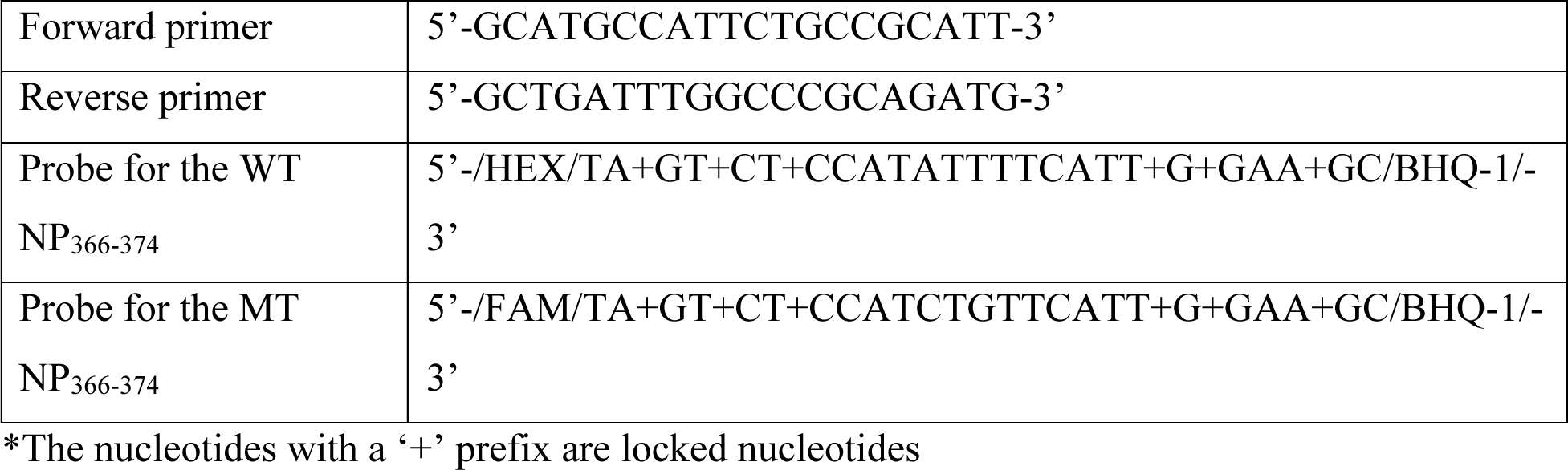

### Surface staining and flow cytometry

Mice received 1.5 ug of CD3e-PE/CF594 in 200 uL sterile PBS 5 minutes before euthanasia. The bronchoalveolar lavage (BAL), lungs, mediastinal lymph node (mLN), and spleen were harvested and processed as previously described [44]. Cells were stained with D^b^NP_366_-BV421, D^b^PA_224_-PE, and K^b^-PB1_703_-APC tetramers provided by NIH Tetramer Core and CD4-BV510, CD8a-BV785, CD44-A700, CD62L-BV605, CD69-A488, CD103-PerCP/Cy5.5, and Zombie-NIR. Samples were run on a Fortessa X-20 flow cytometer (BD Biosciences) and the percentages of tetramer-positive cells were calculated following analysis with FlowJo software.

### Bootstrapping

For each immune setting, the bootstrapping on AUC was done by iterating the following procedure:

1. On each time point, of which *n* mice were measured, randomly sample *n* mice with replacement. Therefore, each sampled mouse will give one WT and one MT viral load.
2. Calculate the mean of log-transformed WT and MT viral loads, and then compute the AUCs of WT and MT with the means.

This procedure was repeated for a million times, and the bootstrapped AUCs were used to approximate the sampling distribution.

### Sensitivity Analysis

To investigate the effects of link function on the estimates of selection coefficient in transmission, we also conducted the AUC and bootstrapping analysis on linear and Hill function, besides logarithm. Formally,

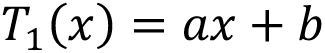

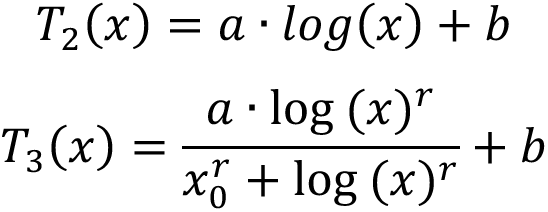

The constant (*b*) is assumed to be 0. In *T*_1_, this assumption indicates transmission does not happen when there is no virus. In *T*_2_ and *T*_3_, it means transmission does not happen when the viral load is below 1 copy/ng, which is the detection limit of our ddPCR system. We notice that the choice of coefficient (*a*) does not affect the estimate of selection coefficient; however, the rate (*r*) and midpoint (*x*_0_) of Hill function will. In **Table S1** we adopted the estimates from Handel et al. [45], where *r* = 4.8 and *x*_0_ = 2.6, and we ran sensitivity analyses on these two parameters.

---

**Fig S1.**
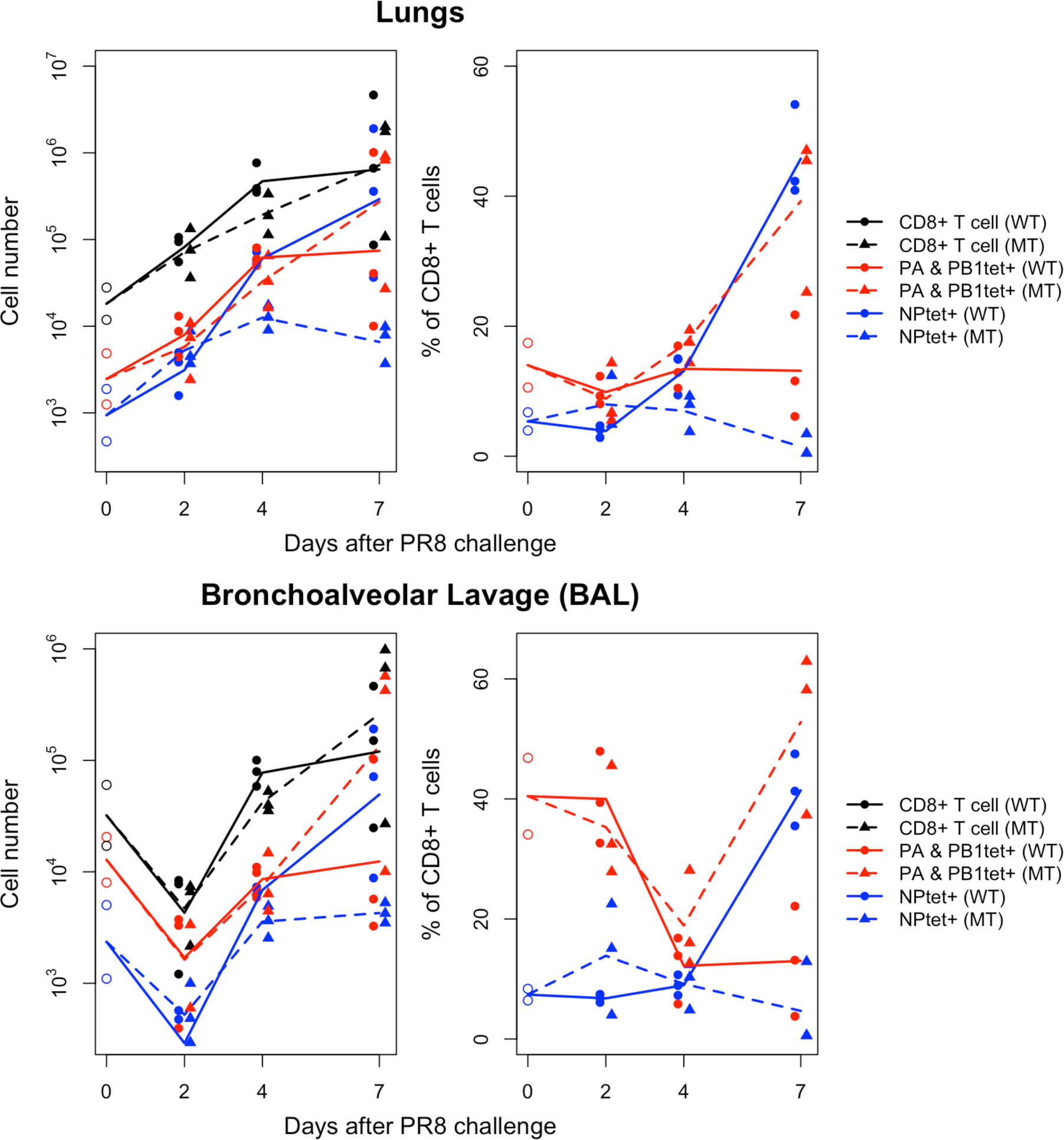
Number and percentage of antigen-specific CD8 T cells in the lung interstitium and airways following challenge with WT or MT viruses. (A, B) i.n. x31-primed B6 mice were rested for 30 days and infected with either WT or MT PR8 viruses. On days 0, 2, 4, and 7 post-challenge, the number and percentage of total CD8+ T cells (black), FluPA- and FluPB1-specific CD8+ T cells (red), and FluNP-specific CD8+ T cells (blue) were assessed in mice challenged with WT (circle) or MT (triangle) viruses.

**Fig S2.**
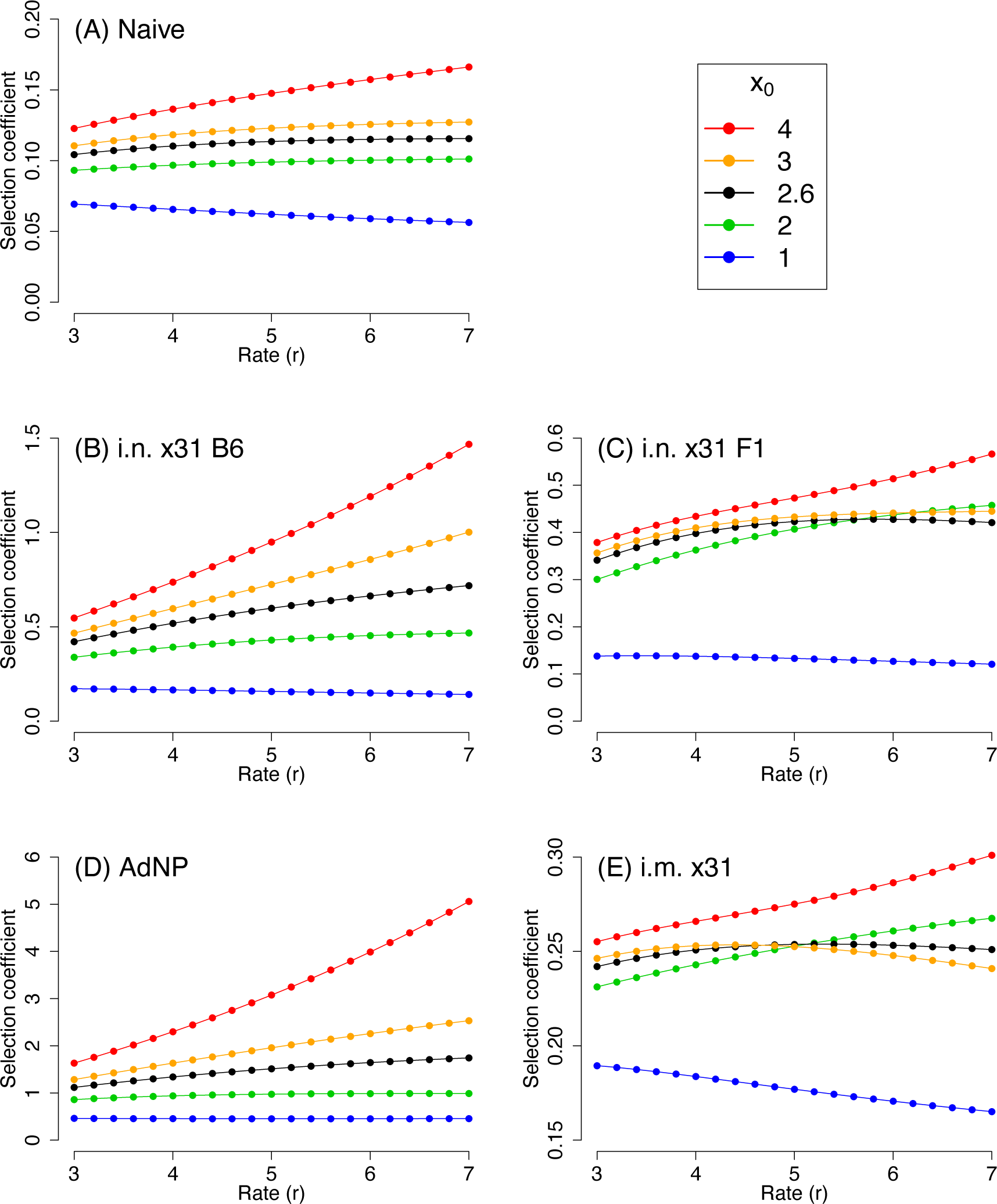
Sensitivity analysis on the rate (*r*) and midpoint (*x*_0_) parameters of Hill function. The selection coefficients were calculated based on the whole infection course (naïve: days 0-9; others: days 0-8).

**Table S1.** Point estimates and 99% empirical bootstrap confidence interval of the selection coefficient of MT under different immune settings and link functions.

| **Table S1** | Point estimates and 99% empirical bootstrap confidence interval of the selection coefficient of MT under different immune settings and link functions. |  |  |
| --- | --- | --- | --- |
|  |  | Link function |  |
| *Days 0-4* | Linear | Logarithmic | Hill |
| Naïve B6<sup>†</sup> | 0.30 [ 0.16, 0.42] | 0.06 [-0.04, 0.12] | 0.08 [-0.06, 0.15] |
| i.n. x31 B6 | 1.31 [ 0.69, 1.71] | 0.20 [ 0.09, 0.30] | 0.55 [ 0.06, 0.77] |
| i.n. x31 F1 | 0.50 [-0.23, 1.01] | 0.17 [-0.02, 0.28] | 0.33 [-0.28, 0.56] |
| AdNP B6 | 7.04 [-2.06, 10.35] | 0.45 [ 0.23, 0.60] | 0.99 [ 0.12, 1.43] |
| i.m. x31 B6 | 0.47 [ 0.28, 0.60] | 0.06 [ 0.03, 0.09] | 0.07 [ 0.01, 0.13] |
| *Days 0-8* |  |  |  |
| Naïve B6<sup>†</sup> | 0.31 [ 0.18, 0.41] | 0.10 [0.003, 0.16] | 0.11 [-0.01, 0.19] |
| i.n. x31 B6 | 1.37 [ 0.79, 1.76] | 0.27 [0.11 , 0.38] | 0.58 [ 0.09, 0.80] |
| i.n. x31 F1 | 0.59 [-0.11, 1.05] | 0.19 [0.01 , 0.31] | 0.42 [-0.27, 0.69] |
| AdNP B6 | 8.39 [-2.90, 12.48] | 0.80 [0.43 , 1.02] | 1.48 [ 0.06, 2.19] |
| i.m. x31 B6 | 0.49 [ 0.23, 0.65] | 0.26 [0.06 , 0.31] | 0.25 [ 0.09, 0.38] |
| <sup>†</sup> For naïve mice, point estimates and bootstrap were conducted over days 0-5 and days 0-9. |  |  |  |

## References

1. Iuliano AD, Roguski KM, Chang HH, Muscatello DJ, Palekar R, Tempia S, et al. Estimates of global seasonal influenza-associated respiratory mortality: a modelling study. Lancet. 2018;391(10127):1285–300. Epub 2017/12/19. doi: 10.1016/S0140-6736(17)33293-2. PubMed PMID: 29248255; PubMed Central PMCID: PMCPMC5935243.

2. Tricco AC, Chit A, Soobiah C, Hallett D, Meier G, Chen MH, et al. Comparing influenza vaccine efficacy against mismatched and matched strains: a systematic review and meta-analysis. BMC Med. 2013;11:153. Epub 2013/06/27. doi: 10.1186/1741-7015-11-153. PubMed PMID: 23800265; PubMed Central PMCID: PMCPMC3706345.

3. Morimoto N, Takeishi K. Change in the efficacy of influenza vaccination after repeated inoculation under antigenic mismatch: A systematic review and meta-analysis. Vaccine. 2018;36(7):949–57. Epub 2018/01/27. doi: 10.1016/j.vaccine.2018.01.023. PubMed PMID: 29373191.

4. Hensley SE, Das SR, Bailey AL, Schmidt LM, Hickman HD, Jayaraman A, et al. Hemagglutinin receptor binding avidity drives influenza A virus antigenic drift. Science. 2009;326(5953):734–6. Epub 2009/11/11. doi: 10.1126/science.1178258. PubMed PMID: 19900932; PubMed Central PMCID: PMCPMC2784927.

5. Lee JM, Eguia R, Zost SJ, Choudhary S, Wilson PC, Bedford T, et al. Mapping person-to-person variation in viral mutations that escape polyclonal serum targeting influenza hemagglutinin. Elife. 2019;8. Epub 2019/08/28. doi: 10.7554/eLife.49324. PubMed PMID: 31452511; PubMed Central PMCID: PMCPMC6711711.

6. Krammer F, Palese P. Advances in the development of influenza virus vaccines. Nat Rev Drug Discov. 2015;14(3):167–82. Epub 2015/02/28. doi: 10.1038/nrd4529. PubMed PMID: 25722244.

7. Soema PC, Kompier R, Amorij JP, Kersten GF. Current and next generation influenza vaccines: Formulation and production strategies. Eur J Pharm Biopharm. 2015;94:251–63. Epub 2015/06/07. doi: 10.1016/j.ejpb.2015.05.023. PubMed PMID: 26047796.

8. Sridhar S. Heterosubtypic T-Cell Immunity to Influenza in Humans: Challenges for Universal T-Cell Influenza Vaccines. Front Immunol. 2016;7:195. Epub 2016/06/01. doi: 10.3389/fimmu.2016.00195. PubMed PMID: 27242800; PubMed Central PMCID: PMCPMC4871858.

9. Paules CI, Marston HD, Eisinger RW, Baltimore D, Fauci AS. The Pathway to a Universal Influenza Vaccine. Immunity. 2017;47(4):599–603. Epub 2017/10/19. doi: 10.1016/j.immuni.2017.09.007. PubMed PMID: 29045889.

10. Clemens EB, van de Sandt C, Wong SS, Wakim LM, Valkenburg SA. Harnessing the Power of T Cells: The Promising Hope for a Universal Influenza Vaccine. Vaccines (Basel). 2018;6(2). Epub 2018/03/29. doi: 10.3390/vaccines6020018. PubMed PMID: 29587436; PubMed Central PMCID: PMCPMC6027237.

11. Sathaliyawala T, Kubota M, Yudanin N, Turner D, Camp P, Thome JJ, et al. Distribution and compartmentalization of human circulating and tissue-resident memory T cell subsets. Immunity. 2013;38(1):187–97. Epub 2012/12/25. doi: 10.1016/j.immuni.2012.09.020. PubMed PMID: 23260195; PubMed Central PMCID: PMCPMC3557604.

12. Farber DL, Yudanin NA, Restifo NP. Human memory T cells: generation, compartmentalization and homeostasis. Nat Rev Immunol. 2014;14(1):24–35. Epub 2013/12/18. doi: 10.1038/nri3567. PubMed PMID: 24336101; PubMed Central PMCID: PMCPMC4032067.

13. Schenkel JM, Masopust D. Tissue-resident memory T cells. Immunity. 2014;41(6):886–97. Epub 2014/12/20. doi: 10.1016/j.immuni.2014.12.007. PubMed PMID: 25526304; PubMed Central PMCID: PMCPMC4276131.

14. O’Neill E, Krauss SL, Riberdy JM, Webster RG, Woodland DL. Heterologous protection against lethal A/HongKong/156/97 (H5N1) influenza virus infection in C57BL/6 mice. J Gen Virol. 2000;81(Pt 11):2689–96. Epub 2000/10/20. doi: 10.1099/0022-1317-81-11-2689. PubMed PMID: 11038381.

15. Kreijtz JH, Bodewes R, van den Brand JM, de Mutsert G, Baas C, van Amerongen G, et al. Infection of mice with a human influenza A/H3N2 virus induces protective immunity against lethal infection with influenza A/H5N1 virus. Vaccine. 2009;27(36):4983–9. Epub 2009/06/23. doi: 10.1016/j.vaccine.2009.05.079. PubMed PMID: 19538996.

16. Wu T, Hu Y, Lee YT, Bouchard KR, Benechet A, Khanna K, et al. Lung-resident memory CD8 T cells (TRM) are indispensable for optimal cross-protection against pulmonary virus infection. J Leukoc Biol. 2014;95(2):215–24. Epub 2013/09/06. doi: 10.1189/jlb.0313180. PubMed PMID: 24006506; PubMed Central PMCID: PMCPMC3896663.

17. McMaster SR, Gabbard JD, Koutsonanos DG, Compans RW, Tripp RA, Tompkins SM, et al. Memory T cells generated by prior exposure to influenza cross react with the novel H7N9 influenza virus and confer protective heterosubtypic immunity. PLoS One. 2015;10(2):e0115725. Epub 2015/02/12. doi: 10.1371/journal.pone.0115725. PubMed PMID: 25671696; PubMed Central PMCID: PMCPMC4324938.

18. Zens KD, Chen JK, Farber DL. Vaccine-generated lung tissue-resident memory T cells provide heterosubtypic protection to influenza infection. JCI Insight. 2016;1(10). Epub 2016/07/29. doi: 10.1172/jci.insight.85832. PubMed PMID: 27468427; PubMed Central PMCID: PMCPMC4959801.

19. Sridhar S, Begom S, Bermingham A, Hoschler K, Adamson W, Carman W, et al. Cellular immune correlates of protection against symptomatic pandemic influenza. Nat Med. 2013;19(10):1305–12. Epub 2013/09/24. doi: 10.1038/nm.3350. PubMed PMID: 24056771.

20. Liang S, Mozdzanowska K, Palladino G, Gerhard W. Heterosubtypic immunity to influenza type A virus in mice. Effector mechanisms and their longevity. J Immunol. 1994;152(4):1653–61. Epub 1994/02/15. PubMed PMID: 8120375.

21. Li ZT, Zarnitsyna VI, Lowen AC, Weissman D, Koelle K, Kohlmeier JE, et al. Why Are CD8 T Cell Epitopes of Human Influenza A Virus Conserved? J Virol. 2019;93(6). Epub 2019/01/11. doi: 10.1128/JVI.01534-18. PubMed PMID: 30626684; PubMed Central PMCID: PMCPMC6401462.

22. Berkhoff EG, de Wit E, Geelhoed-Mieras MM, Boon AC, Symons J, Fouchier RA, et al. Functional constraints of influenza A virus epitopes limit escape from cytotoxic T lymphocytes. J Virol. 2005;79(17):11239–46. Epub 2005/08/17. doi: 10.1128/JVI.79.17.11239-11246.2005. PubMed PMID: 16103176; PubMed Central PMCID: PMCPMC1193597.

23. Berkhoff EG, de Wit E, Geelhoed-Mieras MM, Boon AC, Symons J, Fouchier RA, et al. Fitness costs limit escape from cytotoxic T lymphocytes by influenza A viruses. Vaccine. 2006;24(44-46):6594–6. Epub 2006/07/14. doi: 10.1016/j.vaccine.2006.05.051. PubMed PMID: 16837112.

24. Rimmelzwaan GF, Kreijtz JH, Bodewes R, Fouchier RA, Osterhaus AD. Influenza virus CTL epitopes, remarkably conserved and remarkably variable. Vaccine. 2009;27(45):6363–5. Epub 2009/10/21. doi: 10.1016/j.vaccine.2009.01.016. PubMed PMID: 19840674.

25. Gong LI, Bloom JD. Epistatically interacting substitutions are enriched during adaptive protein evolution. PLoS Genet. 2014;10(5):e1004328. Epub 2014/05/09. doi: 10.1371/journal.pgen.1004328. PubMed PMID: 24811236; PubMed Central PMCID: PMCPMC4014419.

26. Gong LI, Suchard MA, Bloom JD. Stability-mediated epistasis constrains the evolution of an influenza protein. Elife. 2013;2:e00631. Epub 2013/05/18. doi: 10.7554/eLife.00631. PubMed PMID: 23682315; PubMed Central PMCID: PMCPMC3654441.

27. Price DA, Goulder PJ, Klenerman P, Sewell AK, Easterbrook PJ, Troop M, et al. Positive selection of HIV-1 cytotoxic T lymphocyte escape variants during primary infection. Proc Natl Acad Sci U S A. 1997;94(5):1890–5. Epub 1997/03/04. doi: 10.1073/pnas.94.5.1890. PubMed PMID: 9050875; PubMed Central PMCID: PMCPMC20013.

28. Allen TM, Altfeld M, Geer SC, Kalife ET, Moore C, O’Sullivan K M, et al. Selective escape from CD8+ T-cell responses represents a major driving force of human immunodeficiency virus type 1 (HIV-1) sequence diversity and reveals constraints on HIV-1 evolution. J Virol. 2005;79(21):13239–49. Epub 2005/10/18. doi: 10.1128/JVI.79.21.13239-13249.2005. PubMed PMID: 16227247; PubMed Central PMCID: PMCPMC1262562.

29. Bowen DG, Walker CM. Mutational escape from CD8+ T cell immunity: HCV evolution, from chimpanzees to man. J Exp Med. 2005;201(11):1709–14. Epub 2005/06/09. doi: 10.1084/jem.20050808. PubMed PMID: 15939787; PubMed Central PMCID: PMCPMC2213256.

30. Valkenburg SA, Quinones-Parra S, Gras S, Komadina N, McVernon J, Wang Z, et al. Acute emergence and reversion of influenza A virus quasispecies within CD8+ T cell antigenic peptides. Nat Commun. 2013;4:2663. Epub 2013/11/01. doi: 10.1038/ncomms3663. PubMed PMID: 24173108.

31. Thomas PG, Keating R, Hulse-Post DJ, Doherty PC. Cell-mediated protection in influenza infection. Emerg Infect Dis. 2006;12(1):48–54. Epub 2006/02/24. doi: 10.3201/eid1201.051237. PubMed PMID: 16494717; PubMed Central PMCID: PMCPMC3291410.

32. Wu T, Guan J, Handel A, Tscharke DC, Sidney J, Sette A, et al. Quantification of epitope abundance reveals the effect of direct and cross-presentation on influenza CTL responses. Nat Commun. 2019;10(1):2846. Epub 2019/06/30. doi: 10.1038/s41467-019-10661-8. PubMed PMID: 31253788; PubMed Central PMCID: PMCPMC6599079.

33. Webby RJ, Andreansky S, Stambas J, Rehg JE, Webster RG, Doherty PC, et al. Protection and compensation in the influenza virus-specific CD8+ T cell response. Proc Natl Acad Sci U S A. 2003;100(12):7235–40. Epub 2003/05/31. doi: 10.1073/pnas.1232449100. PubMed PMID: 12775762; PubMed Central PMCID: PMCPMC165859.

34. Smith AP, Moquin DJ, Bernhauerova V, Smith AM. Influenza Virus Infection Model With Density Dependence Supports Biphasic Viral Decay. Front Microbiol. 2018;9:1554. Epub 2018/07/26. doi: 10.3389/fmicb.2018.01554. PubMed PMID: 30042759; PubMed Central PMCID: PMCPMC6048257.

35. Uddback IE, Pedersen LM, Pedersen SR, Steffensen MA, Holst PJ, Thomsen AR, et al. Combined local and systemic immunization is essential for durable T-cell mediated heterosubtypic immunity against influenza A virus. Sci Rep. 2016;6:20137. Epub 2016/02/03. doi: 10.1038/srep20137. PubMed PMID: 26831578; PubMed Central PMCID: PMCPMC4735591.

36. Vitiello A, Yuan L, Chesnut RW, Sidney J, Southwood S, Farness P, et al. Immunodominance analysis of CTL responses to influenza PR8 virus reveals two new dominant and subdominant Kb-restricted epitopes. J Immunol. 1996;157(12):5555–62. Epub 1996/12/15. PubMed PMID: 8955206.

37. Flynn KJ, Belz GT, Altman JD, Ahmed R, Woodland DL, Doherty PC. Virus-specific CD8+ T cells in primary and secondary influenza pneumonia. Immunity. 1998;8(6):683–91. Epub 1998/07/09. doi: 10.1016/s1074-7613(00)80573-7. PubMed PMID: 9655482.

38. Crowe SR, Miller SC, Shenyo RM, Woodland DL. Vaccination with an acidic polymerase epitope of influenza virus elicits a potent antiviral T cell response but delayed clearance of an influenza virus challenge. J Immunol. 2005;174(2):696–701. Epub 2005/01/07. doi: 10.4049/jimmunol.174.2.696. PubMed PMID: 15634888.

39. Slutter B, Van Braeckel-Budimir N, Abboud G, Varga SM, Salek-Ardakani S, Harty JT. Dynamics of influenza-induced lung-resident memory T cells underlie waning heterosubtypic immunity. Sci Immunol. 2017;2(7). Epub 2017/08/08. doi: 10.1126/sciimmunol.aag2031. PubMed PMID: 28783666; PubMed Central PMCID: PMCPMC5590757.

40. Pizzolla A, Wakim LM. Memory T Cell Dynamics in the Lung during Influenza Virus Infection. J Immunol. 2019;202(2):374–81. Epub 2019/01/09. doi: 10.4049/jimmunol.1800979. PubMed PMID: 30617119.

41. Kucharski AJ, Lessler J, Read JM, Zhu H, Jiang CQ, Guan Y, et al. Estimating the life course of influenza A(H3N2) antibody responses from cross-sectional data. PLoS Biol. 2015;13(3):e1002082. Epub 2015/03/04. doi: 10.1371/journal.pbio.1002082. PubMed PMID: 25734701; PubMed Central PMCID: PMCPMC4348415.

42. Calderon BM, Danzy S, Delima GK, Jacobs NT, Ganti K, Hockman MR, et al. Dysregulation of M segment gene expression contributes to influenza A virus host restriction. PLoS Pathog. 2019;15(8):e1007892. Epub 2019/08/16. doi: 10.1371/journal.ppat.1007892. PubMed PMID: 31415678; PubMed Central PMCID: PMCPMC6695095.

43. Davison AC, Hinkley DV. Bootstrap methods and their application. Cambridge; New York, NY, USA: Cambridge University Press; 1997. x, 582 p. p.

44. Wein AN, Dunbar PR, McMaster SR, Li ZT, Denning TL, Kohlmeier JE. IL-36gamma Protects against Severe Influenza Infection by Promoting Lung Alveolar Macrophage Survival and Limiting Viral Replication. J Immunol. 2018;201(2):573–82. Epub 2018/06/01. doi: 10.4049/jimmunol.1701796. PubMed PMID: 29848754; PubMed Central PMCID: PMCPMC6089355.

45. Handel A, Longini IM, Jr., Antia R. Neuraminidase inhibitor resistance in influenza: assessing the danger of its generation and spread. PLoS Comput Biol. 2007;3(12):e240. Epub 2007/12/12. doi: 10.1371/journal.pcbi.0030240. PubMed PMID: 18069885; PubMed Central PMCID: PMCPMC2134965.

